# Cirrhosis severity modulates proteomic and immune landscapes in hepatocellular carcinoma

**DOI:** 10.1101/2025.11.24.690286

**Authors:** Raksha K.R. Shastry, Adithya Kale, Sui Seng Tee

## Abstract

Hepatocellular carcinoma (HCC) most often arises in cirrhotic livers, yet the biological impact of cirrhosis severity on the tumor proteome remains poorly understood. Here, we performed a comprehensive quantitative proteomic analysis of 141 HCC tumors stratified by the degree of cirrhosis to delineate fibrosis-related molecular heterogeneity. Principal component analysis revealed substantial overlap between groups, indicating that increasing cirrhosis severity produces incremental rather than global proteomic shifts. Nevertheless, 295 proteins showed significant differential abundance (*Hedge’s g > 0.3, p < 0.05*), including upregulation of HMOX1 and LRSAM1 and downregulation of CYP27A1 in Medium/High cirrhosis. Pathway enrichment highlighted alterations in organelle assembly, heme metabolism, and extracellular-matrix disassembly, alongside reduced cytoskeletal and focal-adhesion integrity. Proteome-based immune deconvolution suggested proportional remodeling of natural-killer and B-cell subsets with advancing cirrhosis. A five-gene signature derived from the most upregulated proteins (*HMOX1, LRSAM1, MAP2, EDEM3, RFC3*) was associated with significantly worse overall survival in the TCGA-LIHC cohort (*Hazard ratio 1.7, p = 0.0033*). Together, these findings demonstrate that cirrhosis severity subtly but measurably reshapes the tumor proteome and immune landscape in HCC, reflecting a continuum of biological remodeling rather than discrete molecular states. Incorporating cirrhosis gradation into molecular classification may improve risk stratification, biomarker discovery, and the design of precision therapies for HCC.

## Introduction

Hepatocellular carcinoma (HCC) accounts for 85-90% of all primary liver cancers, with cirrhosis as the main underlying risk factor in 85% cases [1,2]. The transition from chronic hepatic injury to cirrhosis is shaped by persistent inflammation, progressive fibrosis, and cycles of cellular regeneration, which fundamentally alter the hepatic environment and facilitate the malignant transformation of hepatocytes [3]. Traditionally, cirrhosis has been regarded as a uniform high-risk state for HCC development; however, emerging evidence underscores the heterogeneity within cirrhotic disease in both clinical outcomes and molecular behavior [4].

Key mechanisms, such as chronic inflammation, accrual of somatic mutations, oxidative stress, and impaired immune surveillance, are inextricably linked to the degree of underlying liver fibrosis, influencing the molecular hallmarks and aggressiveness of HCC [5,6]. Despite the longstanding dichotomous clinical division between “cirrhotic” and “non-cirrhotic” HCC, recent reports highlight that tumors arising in non-cirrhotic or minimally cirrhotic livers have a distinct phenotype, including more advanced stage at diagnosis, larger tumor size, and unique etiological risk factors, but may paradoxically demonstrate improved survival in some cohorts [7].

Yet, most comparative studies lump all cirrhotic or non-cirrhotic cases together, failing to parse out the biological impact of cirrhosis severity on tumor phenotype[8]. Notably, in non-alcoholic fatty liver disease (NAFLD), HCC can develop both with and without cirrhosis, and early work shows important differences in tumor characteristics between these groups. However, limited studies have specifically stratified HCC patients by the quantitative degree of cirrhosis, representing a critical knowledge gap [9,10].

The molecular drivers of HCC, including TERT promoter mutations, TP53 and CTNNB1 alterations, are found in both cirrhotic and non-cirrhotic HCC, but it remains unclear how the cirrhotic microenvironment modulates the downstream transcriptomic, epigenomic, and especially proteomic landscapes [11,12]. Clinical strategies, such as surveillance with ultrasonography and serum biomarkers (e.g., AFP), are well established for cirrhotic patients, while detection in non-cirrhotic individuals remains challenging. Newer imaging modalities, including gadoxetic acid-enhanced MRI, have improved sensitivity but predominantly benefit cirrhotic cohorts [1]. Chronic injury from hepatitis B and C viruses, alcohol, or non-alcoholic steatohepatitis(NASH) further complicates the pathological continuum and creates distinct molecular subtypes, yet still most comparative profiling remains binary [13,14].

Part of the difficulty in resolving these nuances relates to limitations in stratification tools. While fibrosis scoring systems such as Ishak or METAVIR are clinically available, inconsistencies in pathology interpretation and the temporal evolution of liver injury hinder their widespread use in research-specific subgrouping [15]. This leaves unresolved whether a graded continuum of cirrhotic change, ranging from minimal to high-grade cirrhosis, yields differences in molecular and clinical tumor profiles.

Consequently, rigorous comparative proteomic studies employing robust group-aware methodologies, as outlined in this investigation, are essential to illuminate how cirrhosis severity shapes HCC biology. Addressing this gap holds promise for new biomarker discovery, improved patient stratification, and the development of precision medicine approaches in HCC.

## Methods

### 2.1 Data source and study population

We analyzed mass spectrometry-based proteomics data from 160 paired hepatocellular carcinoma (HCC) tumor samples stratified by cirrhosis severity (Absent/Low: n=102; Medium/High: n=50) [16]. This data is available in Supplementary Table 1. Following quality control procedures (Section 2.2), 152 samples with 6,512 initially quantified proteins were retained for analysis. Sample exclusions and workflow details are provided in the STROBE diagram.

### 2.2 Mass Spectrometry Quality Control

Proteome data quality was assessed using a two-component mixture model evaluating peptide identification counts and protein identification thresholds. Eight samples failing quality thresholds were excluded, ensuring robust proteomic measurements for downstream analysis.

#### Strobe Diagram

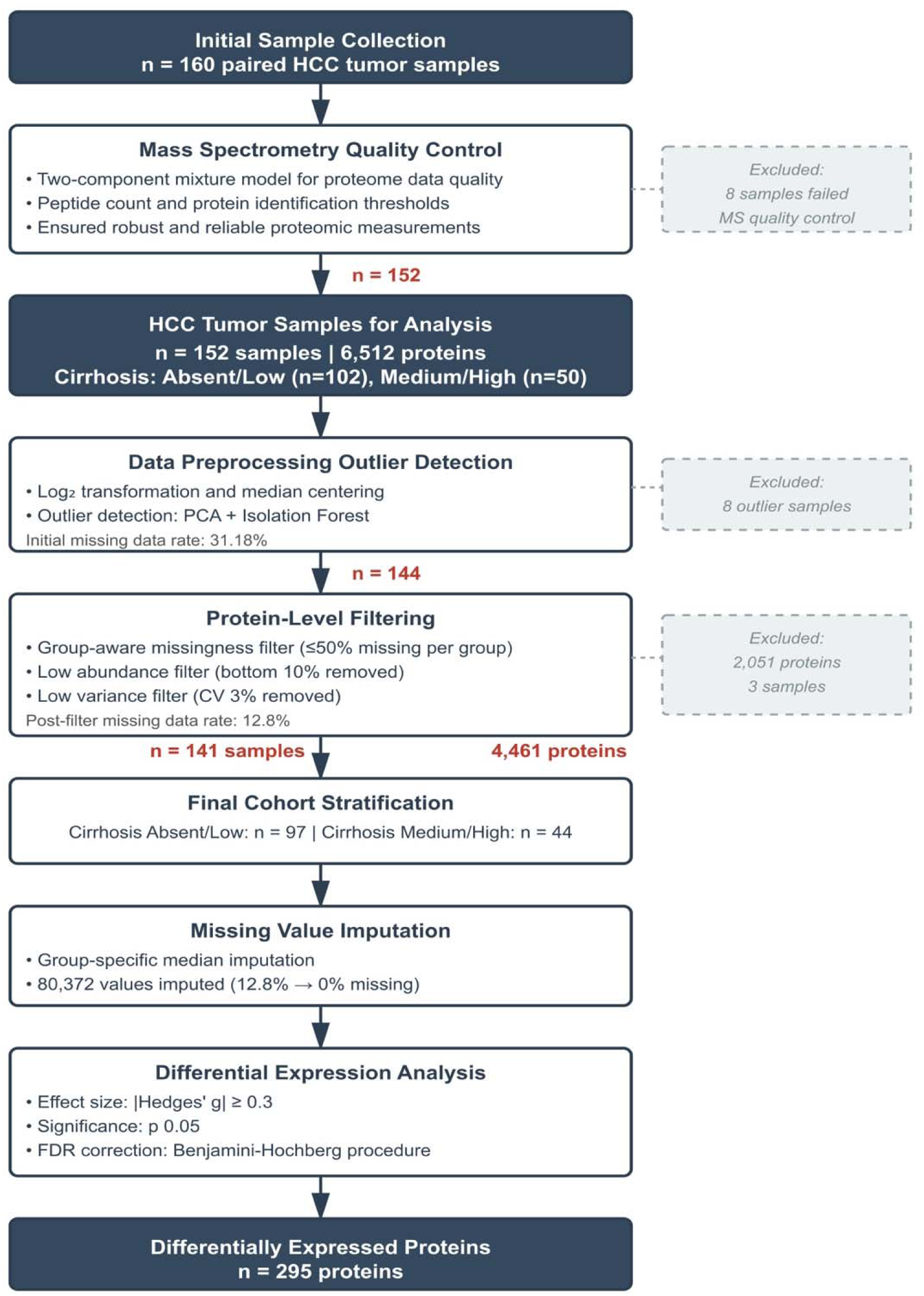

### 2.3 Data preprocessing and normalization

Raw protein intensities underwent log□ transformation to stabilize variance and approximate normal distribution. Sample-wise median centering was applied to correct for systematic technical variation between runs. Outlier samples were identified using principal component analysis (PCA) on principal components combined with machine learning-based isolation forest algorithms. Eight samples exceeding the 95th percentile of anomaly scores were excluded, yielding 144 samples for subsequent filtering.

### 2.4 Protein filtering

To retain high-quality, informative proteins, we applied sequential filters: (1) group-aware missingness filter (proteins with >50% missing values within either cirrhosis group were removed); (2) low abundance filter (bottom 10% by median intensity removed); (3) low variance filter (coefficient of variation <3% removed). This reduced the dataset from 6,512 to 4,461 proteins across 141 samples (3 additional samples were removed due to excessive missingness). Post-filter missing data rate was 12.8%. This data is available in Supplementary Table 2.

### 2.5 Missing value imputation

Missing protein values were imputed separately within each cirrhosis group using group-specific medians to preserve biological differences and prevent cross-group information leakage. This approach imputed 80,372 values, reducing missingness from 12.8% to 0% in the final dataset of 141 samples × 4,461 proteins.

### 2.6 Differential expression analysis

Differential protein expression between cirrhosis groups (Absent/Low: n=97 vs. Medium/High: n=44) was assessed using Hedges’ g effect size with significance threshold p<0.05. False discovery rate correction was applied using the Benjamini-Hochberg procedure. Proteins meeting dual criteria (|Hedges’ g| ≥ 0.3 and FDR-adjusted p<0.05) were considered differentially expressed, yielding 295 proteins. Statistical analyses were performed using Python.

### 2.7 Immune cell deconvolution analysis (CIBERSORTx)

Immune cell composition was estimated from bulk proteomic profiles using CIBERSORTx (v1.0) with the LM22 signature matrix comprising 22 immune cell types. Protein abundance data were first processed using the proteoDeconv R package (v1.0.0): protein identifiers were mapped to HGNC gene symbols, duplicates were collapsed by mean abundance, and missing values were imputed using k-nearest neighbors (k=5). Log -transformed abundances were converted to linear scale and normalized to transcripts per million (TPM)-equivalent units using the convert_to_tpm() function. CIBERSORTx was run in absolute mode without quantile normalization (recommended for proteomic data) with 1,000 permutations for significance testing.

### 2.8 Survival Analysis Using Gene Expression Signature

The five most upregulated proteins in medium/high cirrhosis samples (HMOX1, LRSAM1, MAP2, EDEM3, RFC3; ranked by Hedges’ g) were used to derive a prognostic signature. Corresponding gene expression data from The Cancer Genome Atlas Liver Hepatocellular Carcinoma (TCGA-LIHC) cohort were obtained via GEPIA2. A composite signature score was calculated as the mean log expression of the five genes. Patients were stratified into high versus low expression groups using the median signature score as cutoff. Survival differences were assessed using Kaplan-Meier curves with log-rank tests and Cox proportional hazards regression using Survival Genie 2.0 and GEPIA2 web tools, which automatically computed hazard ratios (HR), 95% confidence intervals (CI), and p-values.

## 3. Results

### 3.1 Patient Demographics and Clinical Characteristics

Table 1 summarizes key clinicopathological features of the 141 hepatocellular carcinoma (HCC) patients included in the proteomics analysis cohort. The majority of patients were male (75%) and presented with solitary tumors (83.7%), reflecting typical epidemiology for HCC. Most tumors were of intermediate size (5-10 cm; 47.5%), with a significant proportion at early/localized tumor stages (TNM I–II: 52.5%, BCLC stage A: 39%). Cirrhosis stratification shows a predominance of Absent/Low (69%) and Medium/High (31%) cirrhosis, underscoring the study’s focus on characterizing the spectrum of cirrhosis severity.

**Table 1:**
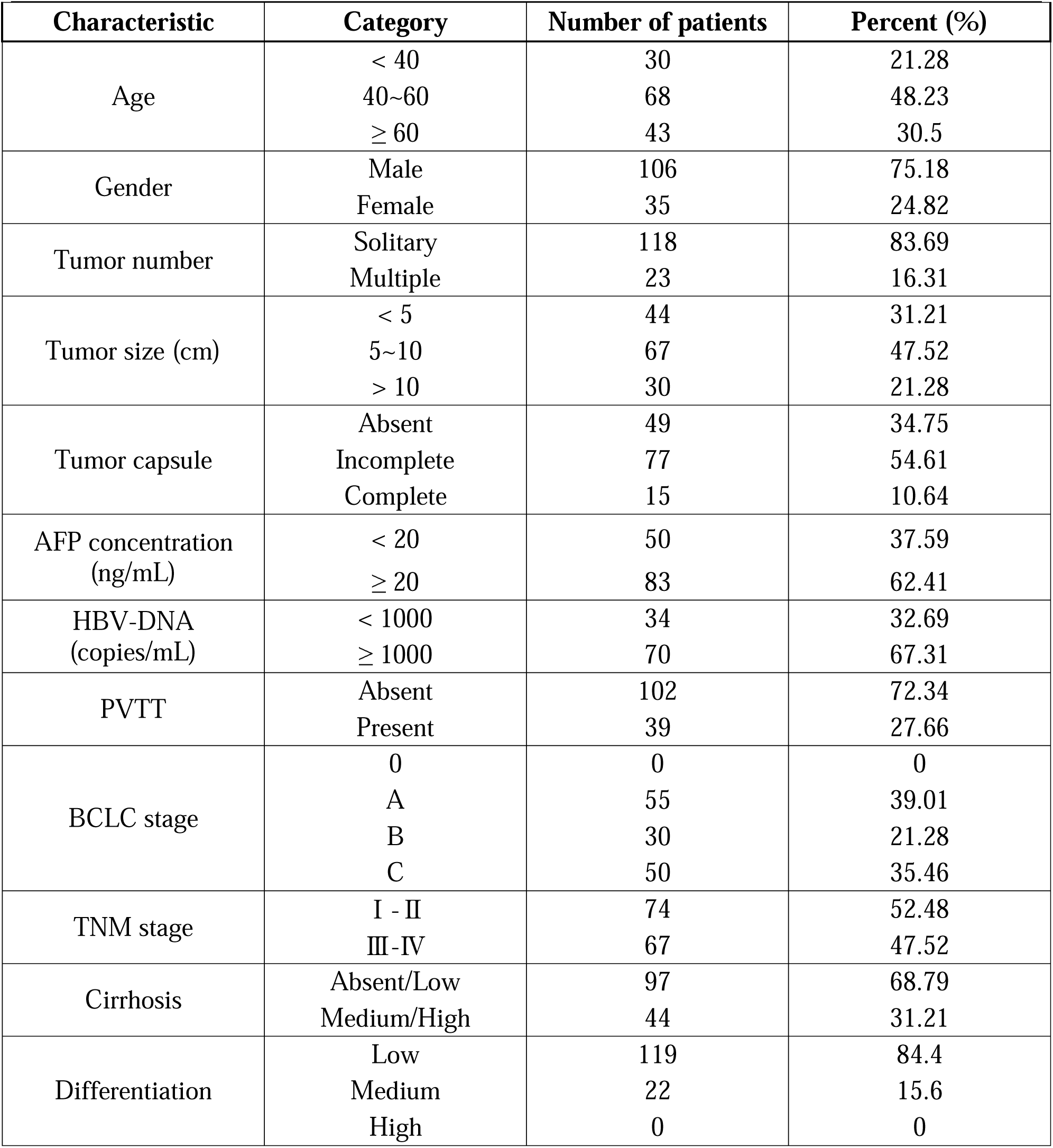
Baseline Clinicopathological Characteristics of the Study Cohort for Proteomics Analysis (n = 141)

Aggressive tumor feature, portal vein tumor thrombus (PVTT present: 27.7%), was moderately observed, highlighting the presence of advanced disease state within the cohort. AFP elevation (≥20 ng/mL) and HBV DNA positivity (≥1000 copies/mL) were present in most cases, consistent with the molecular risk profile prevalent in the region studied.

The balanced representation of cirrhosis levels (Absent/Low, Medium/High) and tumor differentiation grades ensures the proteomic analysis is powered to identify cirrhosis-specific signatures and relate these to clinical and pathological diversity. This clinicopathological breadth allows for nuanced investigation of how proteomic alterations drive disease heterogeneity and progression across cirrhosis microenvironments, with the ultimate goal of informing biomarker discovery and therapeutic strategies for aggressive HCC.

### 3.2 Clustering and Differential Protein Expression

Principal Component Analysis (PCA) was employed to examine the global structure and clustering of proteomic profiles across the study cohort. As visualized in Figure 2A, the top two principal components (PC1 and PC2) accounted for 16% of the total variance in the data. Samples were projected in the PC1-PC2 space and annotated by cirrhosis severity (Absent/Low vs. Medium/High), gender, and year of surgery. There was substantial overlap in PCA coordinates among the cirrhosis groups, as well as across gender and temporal strata, indicating no strong unsupervised clustering driven by these clinical features. These findings suggest that the main sources of variance in the proteomic landscape are not dominated by cirrhosis status, gender, or surgery year.

**Figure 2.**
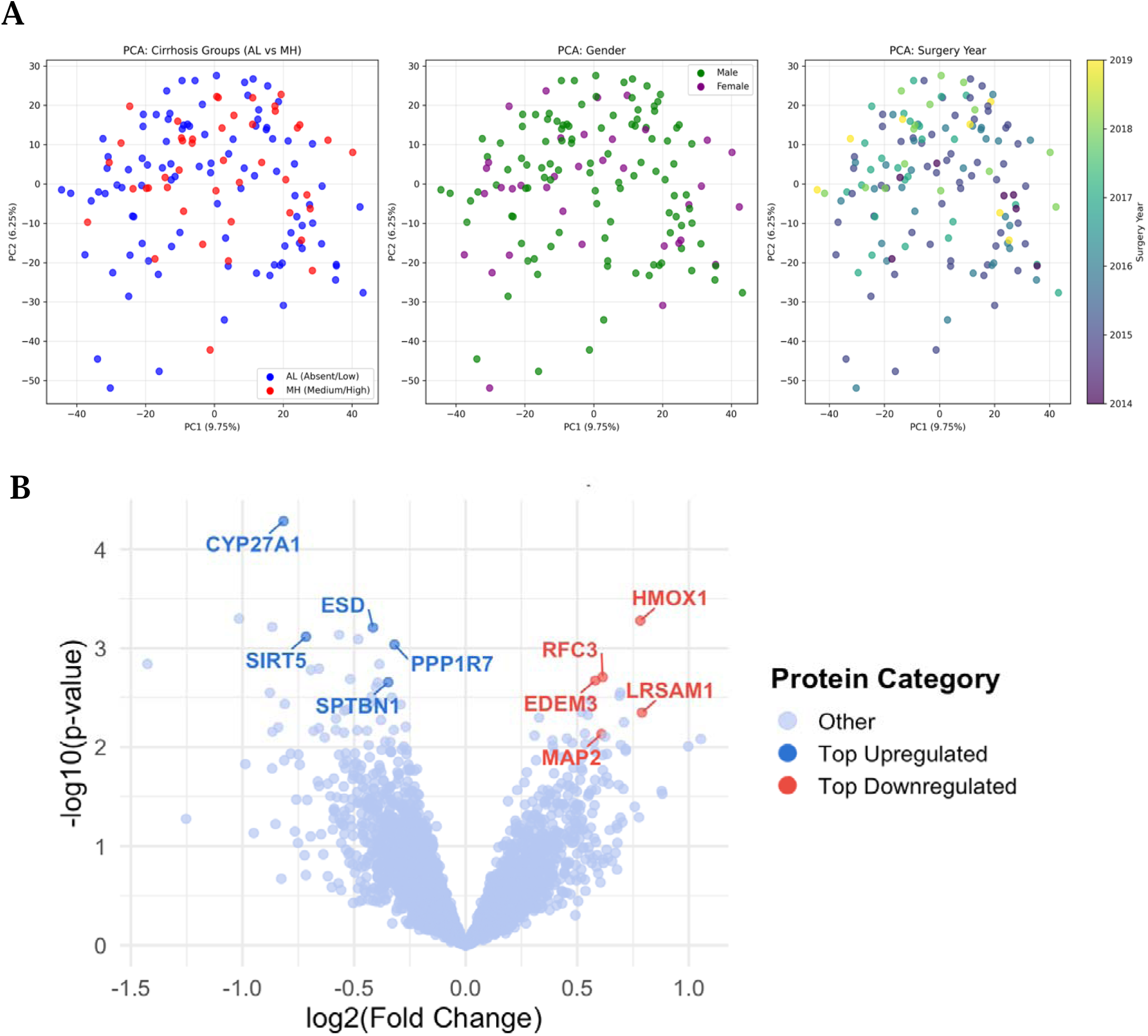
Multi-panel representation of proteomic changes in hepatocellular carcinoma (HCC) according to cirrhosis severity. **(A)** Principal Component Analysis (PCA) plots display the distribution of samples projected onto the first two principal components and annotated by cirrhosis group, gender, and year of surgery, revealing substantial overlap and no distinct clustering by clinical features. **(B)** Volcano plot visualizes the magnitude and significance of differential protein abundance between Absent/Low and Medium/High cirrhosis groups. The top five upregulated (red) and downregulated (blue) proteins are highlighted and labeled. Together, these analyses demonstrate subtle but statistically significant changes in specific proteins despite the absence of marked separation in global proteomic profiles.

To quantitatively evaluate group differences, we performed differential expression analysis between Absent/Low vs. Medium/High cirrhosis groups. Effect sizes for differential abundance were measured using Hedge’s g, and significance was determined with p-values corrected for multiple comparisons (FDR). In total, 295 proteins demonstrated both an absolute Hedge’s g > 0.3 and nominal significance (p < 0.05), including 103 upregulated and 192 downregulated in Medium/High cirrhosis group. The range of effect sizes observed was from -0.64 to 0.73, reflecting mild to moderate disruptions in protein abundance between groups. Representative examples include HMOX1 (Hedge’s g = 0.73, upregulated in Medium/High, p = 0.0005), LRSAM1 (g = 0.61), as well as CYP27A1 (g = -0.64, downregulated in Medium/High, p = 5.2e-5). Notably, many proteins showed more modest but consistent shifts, without exceptionally large effect modifications, aligning with the absence of marked group separation on the unsupervised PCA plots.

Figure 2B highlights the differential protein expression between Absent/Low and Medium/High cirrhosis groups. Each point represents a protein, with its position reflecting both the magnitude of change (log fold change) and statistical significance (–log□□ p-value) [available in Supplementary Table 3]. Consistent with the findings from PCA, which showed substantial overlap of global proteomic profiles without unsupervised clustering by cirrhosis group, the volcano plot reveals that differential abundance between Absent/Low and Medium/High cirrhosis is driven by moderate effect sizes for a subset of proteins rather than dramatic shifts in the global proteomic landscape.

While the majority of proteins demonstrated small or insignificant changes, several proteins exhibited statistically significant differential expression. Notably, HMOX1 and LRSAM1 were strongly upregulated in the Medium/High group, while CYP27A1 and SIRT5 stood out as markedly downregulated. These patterns reinforce the subtle, multifactorial nature of proteomic alterations observed in PCA, indicating that cirrhosis severity imparts moderate but biologically meaningful changes in specific protein pathways rather than global restructuring.

Together, the PCA and volcano plot offer complementary insights: PCA confirms the absence of distinct sample clustering by cirrhosis status, gender, or year, while the volcano plot pinpoints specific proteins with the greatest differential abundance. This integrated view underscores a complex proteomic response to cirrhosis progression, supporting further pathway-focused analysis to elucidate underlying mechanisms.

### 3.3 Pathway Enrichment Analysis

Gene ontology enrichment analysis (Fig. 3) revealed that the most upregulated proteins in Medium/High cirrhosis group were significantly associated with pathways involved in organelle assembly, heme metabolism, nuclear transport, and extracellular matrix disassembly. In contrast, downregulated proteins in Medium/High cirrhosis group showed enrichment for terms related to cytoskeletal structure, focal adhesion, and cell junctions, indicating a loss of cell structural integrity and connectivity with disease progression. Further, both Absent/Low and Medium/High cirrhosis groups displayed distinct patterns of molecular function and cellular component enrichment, including alterations in tubulin and actin binding activities. These findings highlight the specific biological processes and molecular functions disrupted by cirrhosis severity and provide mechanistic insights into the proteomic remodeling underlying hepatocellular carcinoma development.

**Figure 3.**
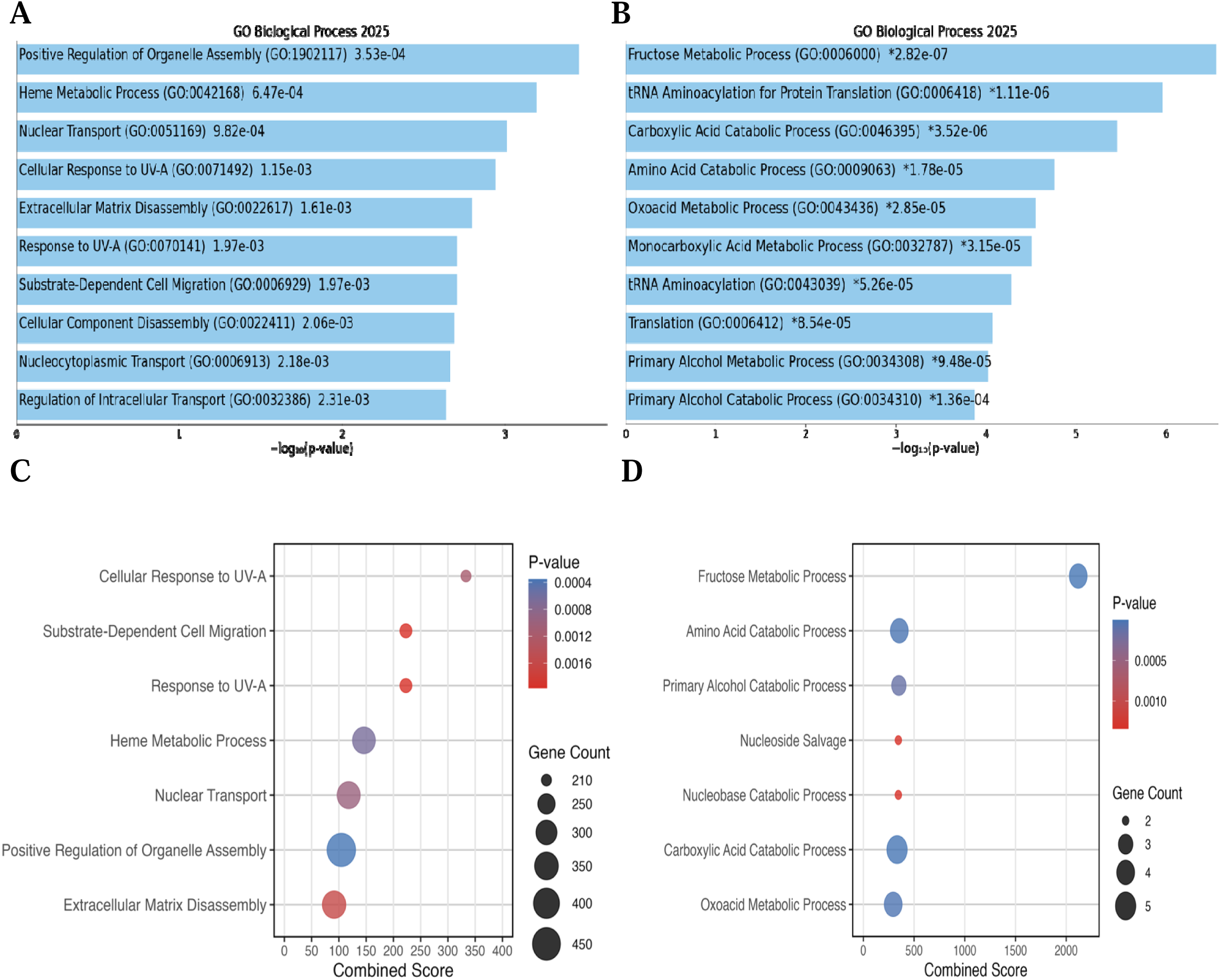
Comparative gene ontology enrichment analysis of upregulated genes in cirrhosis groups. (A) Bar plot showing to enriched GO biological processes for upregulated genes in Medium/High cirrhosis group. (B) Bar plot showing top enriched GO biological processes for upregulated genes in Absent/Low cirrhosis. (C) Dot plot visualizing significant GO biological processes for upregulated genes in Medium/High cirrhosis group; dot size indicates number of genes in each term and color reflects statistical significance (p-value). The x-axis shows the ‘Combined Score,’ which integrates both enrichment significance (p-value) and deviation from expectation (z-score) for each GO term. (D) Dot plot visualizing significant GO biological processes for upregulated genes in Absent/Low cirrhosis; dot size and color are as above. A complete list of enriched terms, including results for Cellular Component and Molecular Function, is provided in Supplementary Figures S1

### 3.4 Deconvolution of immune cell populations by cirrhosis severity

Using proteomic deconvolution and CIBERSORTx, we inferred the relative proportions of 22 immune cell types across all HCC tumor samples (Supplementary Table 4). B cell memory was the most abundant cell population overall, followed by mast cells (resting), T cells CD8, and T cells gamma delta. Visualizing the top immune cell types by mean abundance (Figure 5A, B) revealed broad immune ecosystem diversity in our cohort.

**Figure 5.**
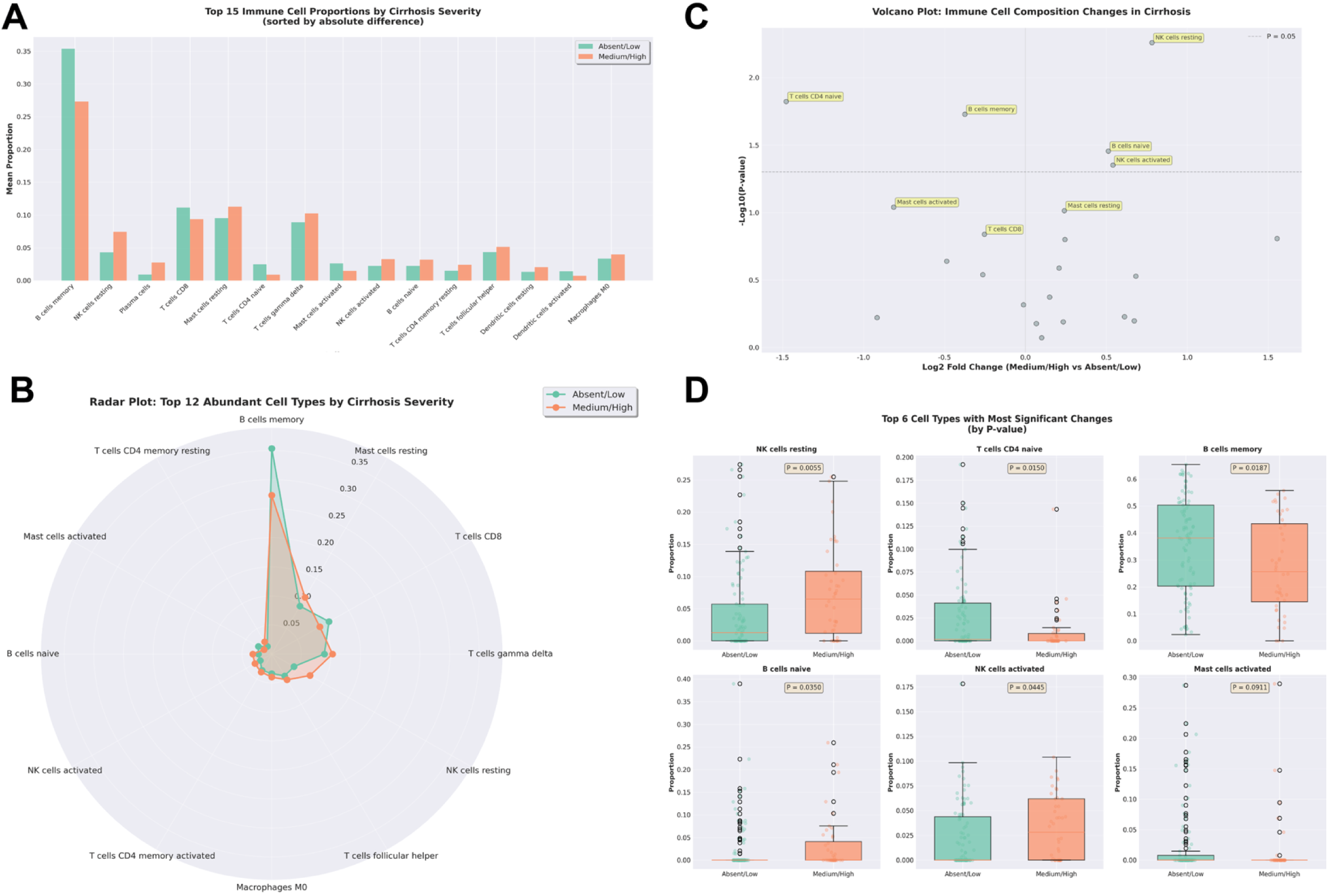
Quantitative immune cell deconvolution of proteomic HCC samples using CIBERSORTx. (A) Bar plot of the top 15 immune cell populations by mean proportion in Absent/Low vs. Medium/High cirrhosis groups (sorted by absolute difference). (B) Radar plot showing the top 12 most abundant immune cell types by group. (C) Volcano plot of immune cell composition differences between cirrhosis groups, plotting log2 fold change (Medium/High vs. Absent/Low) versus nominal significance; highlighted cell types are those with the largest differences. (D) Boxplots of the 6 cell types with the most significant group differences, with p-values indicated from Wilcoxon rank-sum tests (not FDR-adjusted). Together, these panels illustrate prominent immune cell populations in HCC and the proportional redistribution of select immune subsets by cirrhosis severity.

Comparing Absent/Low cirrhosis groups vs. Medium/High, six immune cell types had the largest absolute difference in mean proportions (Figure 5A). NK cells resting, B cells naive, and NK cells activated were increased in Medium/High cirrhosis, while T cells CD4 naive and B cells memory were decreased in Medium/High compared to Absent/Low (see boxplots, Figure 5D). However, after multiple testing correction, none of these changes reached FDR significance (FDR-adjusted p > 0.1), though the nominal p-values for several (NK cells resting: p = 0.006, T cells CD4 naive: p = 0.015, B cells memory: p = 0.019) suggest potential biological relevance. Overall, these results indicate immune compartment remodeling in Medium/High cirrhosis, with proportional shifts most apparent in NK cells and B cell subsets.

### 3.5 Survival analysis of Gene Signature

Patients with high signature expression (n=182) demonstrated significantly reduced overall survival compared to the low expression group (n=182), as shown by the Kaplan-Meier analysis (Figure 6). The hazard ratio was 1.7 (95% CI: 1.2-2.4), with a log-rank p=0.0033, indicating that upregulation of this gene set is associated with increased mortality risk in TCGA-LIHC. These results support the prognostic relevance of this gene signature in cirrhosis-associated HCC.

**Figure 6.**
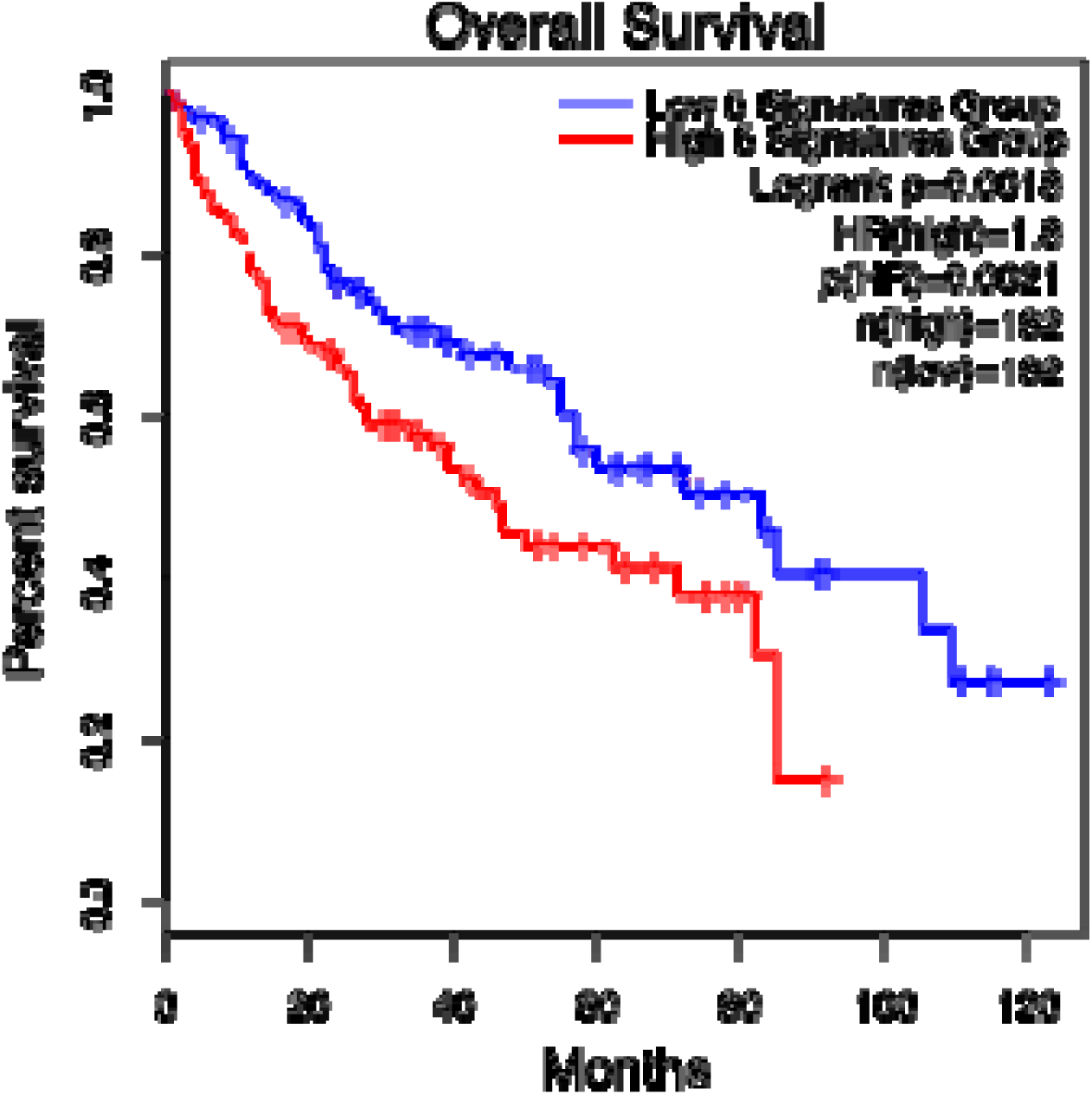
Kaplan-Meier survival analysis of overall survival in TCGA-LIHC stratified by median expression of the five-gene (HMOX1, LRSAM1, MAP2, EDEM3, RFC3) signature. High expression (red, n=185) was associated with significantly worse survival compared to low expression (blue, n=184), HR 1.7 (95% CI: 1.2–2.4), log-rank p=0.0033.

## 4. Discussion

We conducted a comprehensive proteomic analysis of hepatocellular carcinoma (HCC) tumors, carefully stratifying patients by the severity of underlying liver cirrhosis. Our study revealed that, rather than dramatic global shifts, cirrhosis severity mainly drives moderate but meaningful changes in specific protein pathways within HCC. Principal component analysis showed a high degree of overlap among cirrhosis groups, meaning that tumor protein profiles did not cluster distinctly by cirrhosis level. However, we identified 295 proteins with significant differences between Absent/Low and Medium/High cirrhosis, and highlighted key proteins such as HMOX1 and LRSAM1 that are upregulated in Medium/High cirrhosis, while others like CYP27A1 are downregulated.

Pathway enrichment analysis pinpointed biological processes altered by increasing cirrhosis, including organelle assembly, heme metabolism, and cell structural integrity. Deconvolution of the tumor immune microenvironment revealed proportional remodeling of immune cell populations in Medium/High cirrhosis, particularly in NK and B cell subsets, although this did not reach strict statistical significance. Importantly, a gene signature derived from the most upregulated proteins in cirrhosis-associated HCC was linked to significantly worse patient survival, suggesting potential clinical relevance for prognosis and therapy.

Our analysis revealed that the differences in protein abundance across cirrhosis severity in HCC were characterized by moderate effect sizes (Hedges’s g ranging from -0.64 to 0.73), with most proteins showing subtle but statistically significant changes rather than dramatic shifts. Principal component analysis reinforced this, showing substantial overlap between cirrhosis groups without distinct clustering, suggesting that cirrhosis severity produces nuanced rather than categorical proteomic states. This finding supports the concept of HCC as a disease continuum, shaped by incremental alterations within the tumoral microenvironment rather than rigidly defined molecular subtypes[17]. Such gradation has important clinical implications, stratification by cirrhosis severity may better capture the biological complexity seen in HCC and inform precision medicine efforts, moving beyond the dichotomous non-cirrhotic vs. cirrhotic classification that oversimplifies patient heterogeneity [18].

Among the most differentially expressed proteins, HMOX1 (heme oxygenase-1) stood out, being strongly upregulated in tumors arising from Medium/High cirrhotic livers. HMOX1 is central to the oxidative stress response, catalyzing the degradation of heme and mitigating iron-induced cytotoxicity. Its induction in Medium/High cirrhosis aligns with the chronic inflammatory milieu of advanced liver disease, where persistent oxidative insult and elevated heme turnover drive HMOX1 expression as a cytoprotective adaptation. Chronic upregulation of HMOX1 has been linked with increased tumor cell survival and immune evasion, implicating it in HCC progression under oxidative stress [19,20]. In contrast, CYP27A1, an enzyme involved in bile acid synthesis and cholesterol metabolism, was among the most downregulated proteins in Medium/High cirrhosis. Decreased CYP27A1 suggests disturbed bile acid homeostasis and impaired hepatic detoxification, features associated with progressive liver dysfunction and carcinogenesis. Downregulation of CYP27A1 may contribute to altered lipid signaling and exacerbate the fibrotic pro-tumor environment that drives HCC evolution in cirrhosis[21, 22].

Pathway enrichment analysis revealed distinct shifts in biological processes correlating with cirrhosis severity. In Absent/Low cirrhosis, upregulated proteins were strongly associated with focal adhesion, cell junction, and cytoskeletal pathways, maintaining cell structure and connectivity. However, as cirrhosis worsened, gene ontology terms shifted towards organelle assembly, nuclear transport, heme metabolism, and especially extracellular matrix (ECM) disassembly and remodeling (see Fig. 3A-D). This transition suggests that with increasing fibrosis and architectural distortion, HCC tumors undergo proteomic reprogramming that facilitates organelle biogenesis and ECM breakdown, hallmarks of malignant transformation and tumor invasion in the cirrhotic liver. These disruptions reflect the known biology of fibrosis, where hepatocyte-to-stroma interactions, loss of cell polarity, and sustained matrix turnover accelerate carcinogenesis and metastatic potential[23, 24].

Immune deconvolution showed notable, albeit nominally significant, shifts in NK cell and B cell subsets between Absent/Low and Medium/High cirrhosis groups. While these changes did not reach false discovery rate significance, NK cells (resting p=0.006) and naive B cells demonstrated proportional increases in Medium/High cirrhosis, alongside decreases in memory B cells and CD4+ T cells. Even moderate alterations in immune cell composition are biologically meaningful, especially given the critical role of immune surveillance in restraining tumor growth [25]. NK cell expansion may reflect compensatory mechanisms in response to chronic inflammation and persistent viral or metabolic insults [26], while diminished memory B cell fractions suggest impaired adaptive immunity[27]. These trends underscore progressive immune surveillance deterioration in cirrhotic HCC, a process central to tumor evasion and poor clinical outcomes [28].

These findings align with broader HCC immune-profiling studies. Immune deconvolution analyses have reported increases in naive B cells and CD8 T cells in HCC tumor tissue compared to healthy liver, with altered memory B cell populations [29, 30]. However, patterns vary considerably: one CIBERSORT analysis found no significant NK cell differences between HCC and normal tissues [31], contrasting with our observed NK expansion in Medium/High cirrhosis, while DNA methylation–based deconvolution studies reported decreased NK cells in HCC tumors versus adjacent tissue [32], suggesting NK dynamics differ between cirrhotic and malignant states. High-resolution immunohistochemistry studies show all B cell subsets, including naive and memory B cells, are significantly decreased within HCC tumors [30], though atypical memory B cells can acquire immunosuppressive, Breg-like phenotypes in the tumor microenvironment [33]. These divergent findings likely reflect differences in tissue compartments analyzed (e.g., blood vs tumor vs cirrhotic tissue), disease stage, and etiology, highlighting the dynamic nature of immune remodeling during HCC pathogenesis [34].

Our study identified a five-gene signature: HMOX1, LRSAM1, MAP2, EDEM3, and RFC3, whose elevated expression in cirrhosis-associated HCC demonstrated clear prognostic value in the TCGA-LIHC dataset (hazard ratio [HR] = 1.7, p = 0.0033). Patients with high median expression of this gene set had significantly reduced overall survival, indicating that upregulation of these proteins may serve as a powerful marker of poor prognosis in HCC with Medium/High cirrhosis. As molecular profiling becomes increasingly integrated into clinical practice, these proteins present promising avenues for risk stratification, allowing clinicians to identify high-risk patients who may benefit from intensified surveillance or tailored therapies [35]. Importantly, these proteomic markers could complement established biomarkers such as alpha-fetoprotein (AFP), which, while widely used, has limited sensitivity and specificity in certain patient subsets [36]. The combination of classic and novel biomarkers may enhance detection and prognostication, especially for tumors arising in the context of severe cirrhotic remodeling.

The pathway analysis, highlighted by disruptions in organelle assembly, ECM remodeling, and immune microenvironment reprogramming, points to targetable vulnerabilities that may be specific to cirrhosis-driven HCC. For example, the upregulation of proteins involved in heme metabolism (e.g., HMOX1) and nuclear transport suggests that inhibitors targeting these processes could selectively affect tumor cells thriving in fibrotic, inflamed livers[19, 20]. Conversely, pathways related to structural integrity and bile acid metabolism that are lost in Medium/High cirrhosis represent potential areas for restoring hepatic function and impeding carcinogenesis [37]. Notably, immune profiling revealed NK cell and B cell subset changes, suggesting that immunotherapeutic strategies, such as NK cell modulation or adaptive immune enhancement, may need to be tailored according to cirrhosis severity [38]. Such insights underscore the need for precision medicine approaches that factor in cirrhosis-driven proteomic remodeling when designing effective therapies, moving beyond “one-size-fits-all” regimens and towards truly personalized management for HCC patients.

Previous studies have typically approached hepatocellular carcinoma (HCC) as a binary disease arising from cirrhotic versus non-cirrhotic livers, often contrasting clinical features and molecular profiles between these two groups [7, 9, 10] . For example, Gaddikeri et al. (2014) and Rastogi (2020) show that non-cirrhotic HCC frequently presents with larger, solitary tumors and is associated with distinct risk factors, such as hepatitis B virus infection or metabolic syndrome, whereas cirrhotic HCC exhibits a higher incidence of vascular invasion, multifocality, and is predominantly driven by chronic hepatitis C or alcohol abuse[7,10]. Pinyopornpanish et al. (2021) further revealed that non-cirrhotic HCC in NAFLD patients has unique clinical and demographic characteristics compared to cirrhotic HCC, with differences in age distribution, tumor size, and surgical eligibility [9]. However, these studies largely treated cirrhosis as a categorical state and did not consider gradations of fibrosis or their impact on tumor biology.

In contrast, our study demonstrates that even within the realm of cirrhotic HCC, the severity of cirrhosis exerts meaningful influences on the molecular phenotype, generating nuanced proteomic and pathway changes rather than sharp molecular boundaries. This graded approach reveals incremental remodeling in protein pathways, immune cell composition, and gene expression as cirrhosis advances, challenging the paradigm of a binary classification.

Concordance with previous transcriptomic and genomic research comes from studies showing the frequent presence of TP53, CTNNB1, and TERT promoter mutations in HCC irrespective of background liver disease, although prevalence and mutational effects may vary with etiology and geography [11, 12]. For instance, Kumar et al. (2023) and Lombardo et al. (2020) spotlight the role of TP53 and CTNNB1 hotspot mutations in HCC pathogenesis, with poor survival associated with mutation-driven molecular subtypes [11, 12]. Our proteomic findings support these observations by linking molecular signatures to survival outcomes. Nevertheless, our data suggests that cirrhosis severity further stratifies the biological risk and provides a lens to refine the molecular taxonomy of HCC, a perspective not captured by most earlier studies. Thus, while there is major concordance regarding driver mutations and certain pathway disruptions, our work highlights cirrhosis severity as a modulator of tumoral heterogeneity and clinical trajectory, offering new directions for both prognosis and therapeutic targeting in HCC.

This study incorporates several methodological strengths that enhance the reliability and clarity of our findings. Notably, we utilized a group-aware imputation strategy that preserves underlying biological differences between cirrhosis groups, minimizing biases introduced by missing data, a frequent challenge in proteomic profiling. The analytical workflow combined rigorous statistical filtering with strict false discovery rate (FDR) correction, enhancing confidence in the identified protein signatures and enriched pathways. Another key strength is the integration of clinical characteristics, proteomic data, and survival outcomes, which provides a multi-layered perspective on the biological and prognostic relevance of cirrhosis severity in HCC. Importantly, our approach moves beyond the conventional binary cirrhotic/non-cirrhotic frameworks; by stratifying patients according to cirrhosis severity, we could reveal more nuanced proteomic and clinical heterogeneity, thus capturing the full spectrum of disease biology.

However, several limitations should be acknowledged. The cohort size, though adequate for discovery, was moderate overall and especially limited for the highest severity cirrhosis subgroup, which may affect statistical power and restrict the detection of subtle differences. Being a single-center study, generalizability may be compromised, as patient demographics and treatment protocols can vary widely across centers. The predominance of HBV (hepatitis B virus) etiology within our cohort also raises questions about the applicability of our findings to HCCs arising from other causes, such as NASH, alcohol-related liver disease, or HCV infection. While mass spectrometry-based proteomics offers deep coverage, it has inherent limitations in capturing the full range of post-translational modifications and protein isoforms relevant to tumor progression. Lastly, the cross-sectional nature of the study limits causal inference; future longitudinal and multi-etiology cohorts, ideally with validation in external datasets, will be critical to confirm and extend these insights.

To build upon these findings, future research should prioritize validation of the identified proteomic signatures in independent, multi-center cohorts that encompass a broader spectrum of underlying etiologies, including NASH, alcohol-associated, and HCV-related liver disease. Longitudinal studies tracking the proteomic evolution from cirrhosis through the onset and progression of HCC would provide insight into dynamic molecular changes and potentially reveal early biomarkers for transformation. Functional studies of key proteins identified here, such as HMOX1 and CYP27A1, are urgently needed to dissect their mechanistic roles in tumor biology, treatment response, and immune modulation. Additionally, integrating proteomic data with genomic and transcriptomic analyses in a multiomics framework will enable a more comprehensive understanding of HCC pathogenesis and may identify novel avenues for therapeutic intervention and precise risk stratification.

This study addresses the critical gap in existing hepatocellular carcinoma research, the lack of stratification by cirrhosis severity and its biological impact. While prior work has treated cirrhosis as a uniform risk state or compared cirrhotic versus non-cirrhotic HCC in a binary fashion, our findings reveal that the degree of cirrhosis imparts moderate but biologically meaningful changes in tumor proteomic signatures and immune remodeling. By demonstrating that incremental fibrosis yields distinct molecular features within HCC, we highlight how cirrhosis severity modulates the clinical and molecular phenotype, a factor overlooked in most previous studies. These results support a move toward stratified, rather than binary, approaches in both future HCC research and clinical practice, paving the way for improved biomarker discovery, risk assessment, and precision medicine tailored to the nuanced understanding of tumor microenvironments.

## Supplementary Figures

**Figure S1.**
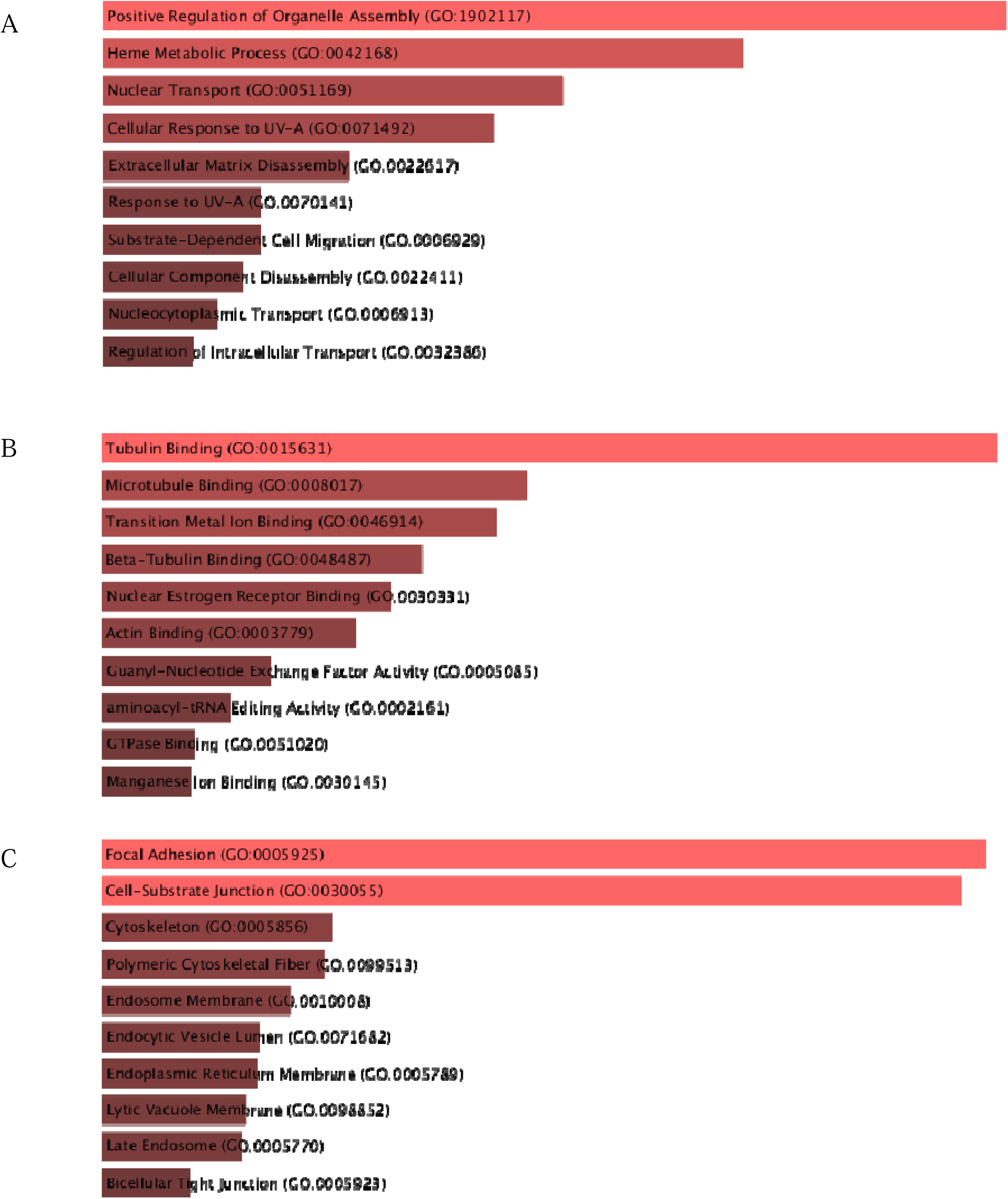
(A-C): Top enriched Gene Ontology (GO) terms for the most upregulated genes in the Medium/High cirrhosis group compared to the Absent/Low cirrhosis group, displayed for: (a) Biological Process, (b) Cellular Component, (c) Molecular Function.

**Fig S2.**
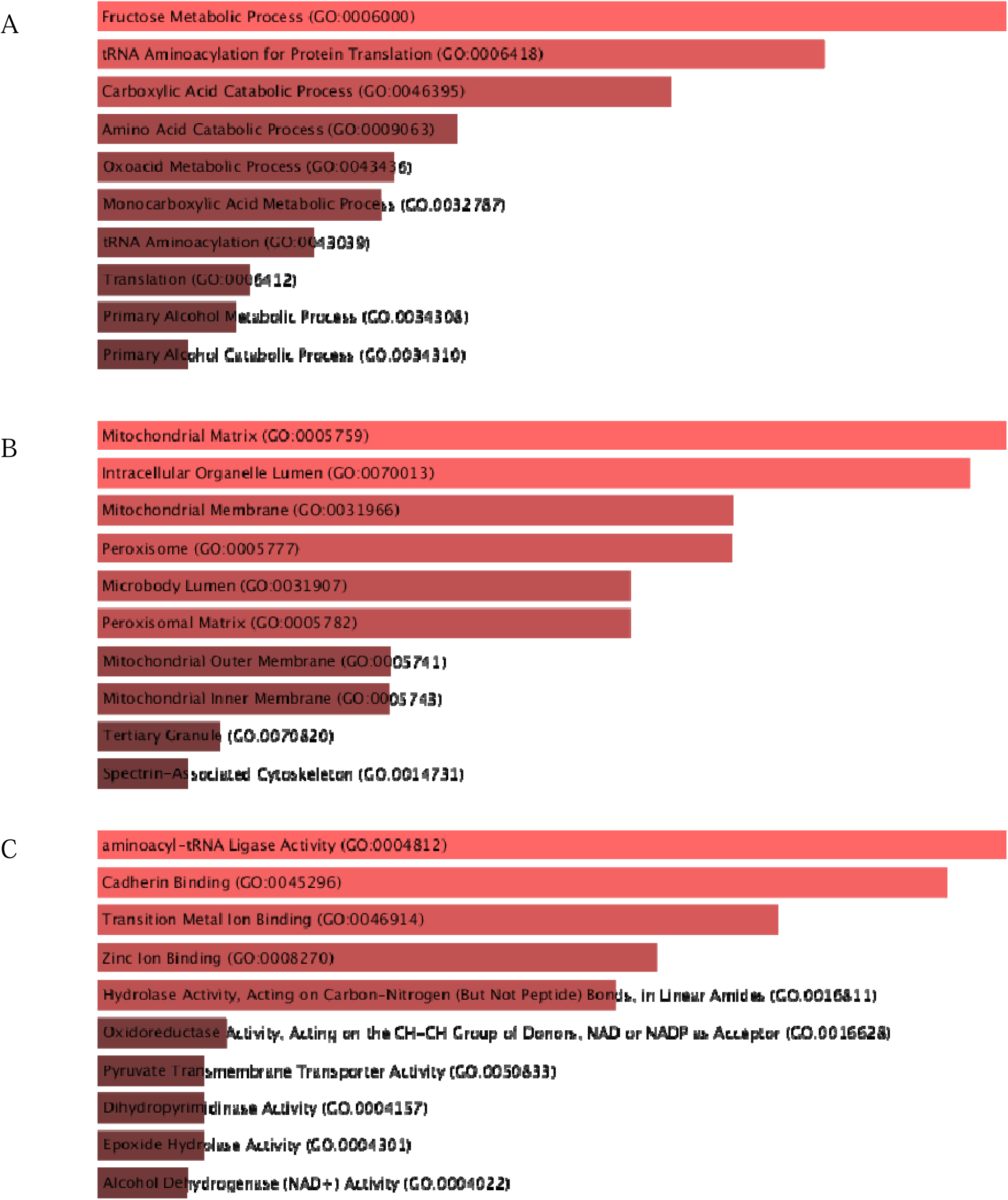
(A-C): Top enriched GO terms for the most downregulated genes in the Medium/High cirrhosis group shown for (d) Biological Process, (e) Cellular Component, (f) Molecular Function.

**Fig S3.**
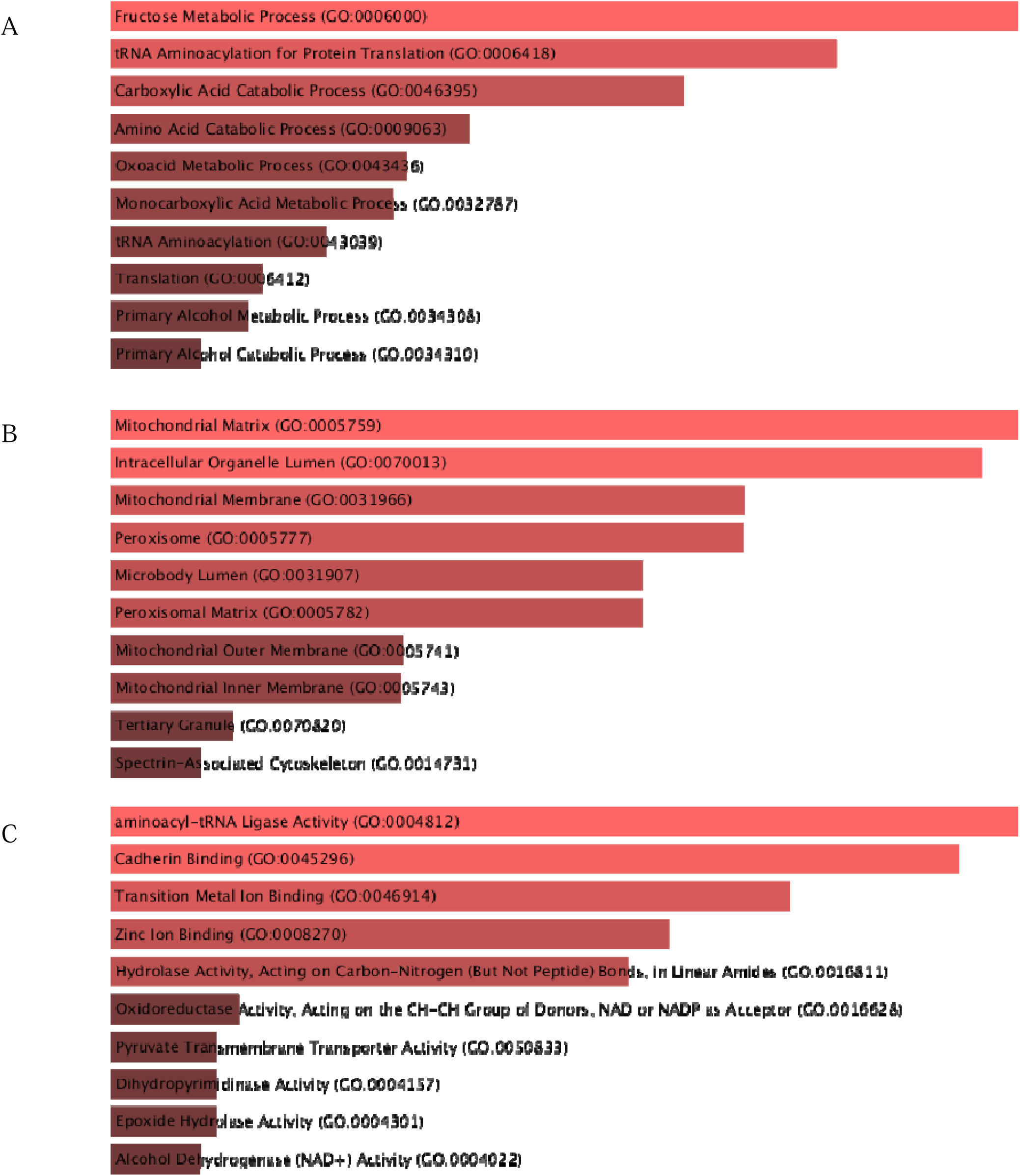
(A-C): Top enriched GO terms for the most upregulated genes in the Absent/Low cirrhosis group shown for (g) Biological Process, (h) Cellular Component, (i) Molecular Function.

**Fig S4.**
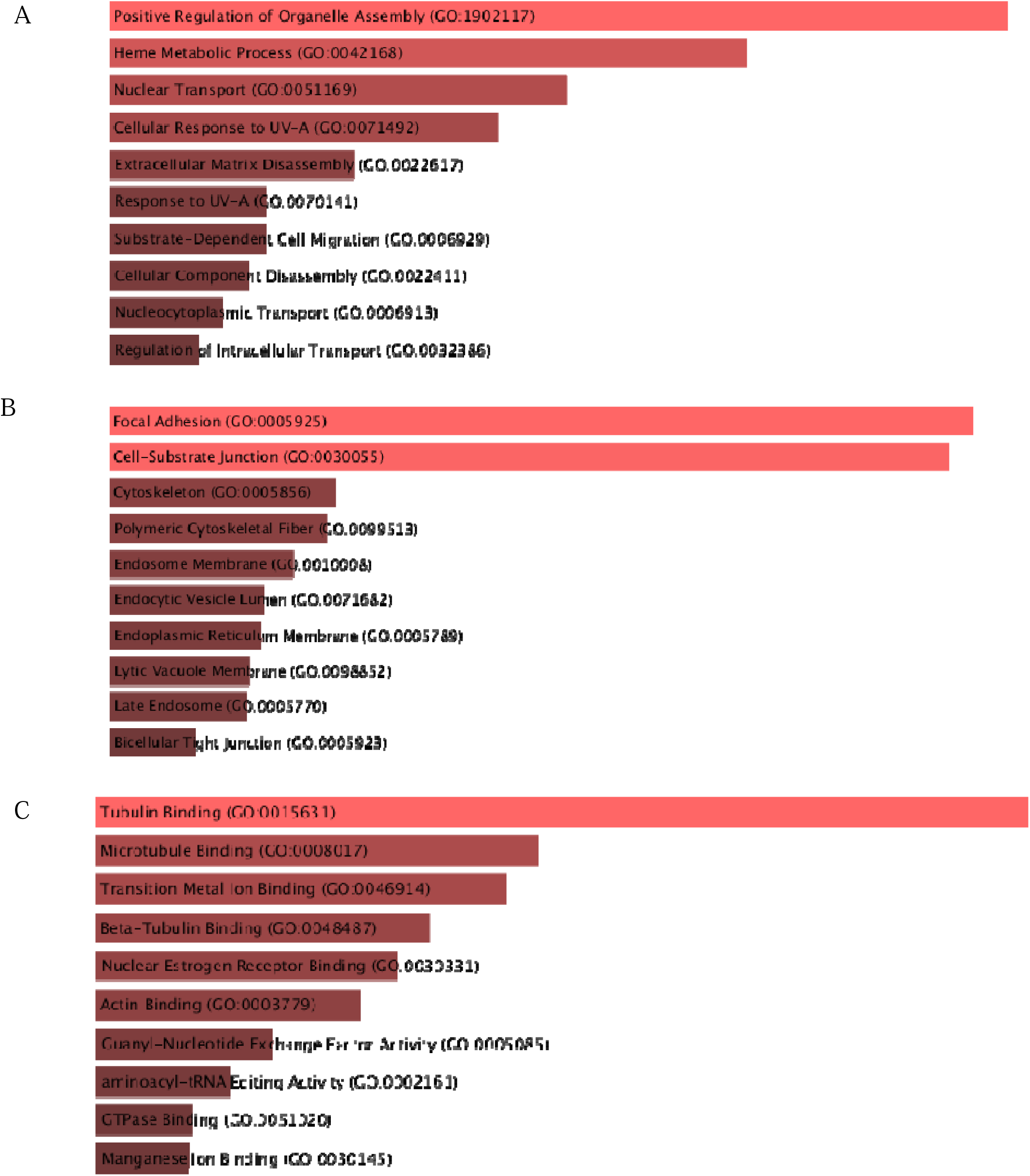
(A-C): Top enriched GO terms for the most downregulated genes in the Absent/Low cirrhosis group shown for (j) Biological Process, (k) Cellular Component, (l) Molecular Function.

